# Single-neuron selectivity rules underlying population representations in prefrontal cortex

**DOI:** 10.1101/2023.01.16.524214

**Authors:** Claudia Böhm, Albert K. Lee

## Abstract

Neurons in prefrontal cortex exhibit selectivity for a range of behaviorally relevant variables, such as sensory stimuli, place, time, choice, context, and category. At the population level, representations of these variables can be organized in various ways—for instance, geometrically or hierarchically. However, how selectivity for each feature is distributed across cells and how this gives rise to such population-level representational structure is unclear. To address these questions, we analyzed coding for space, time, and category in rats performing a navigational task featuring two behavioral categories, starts and goals, each consisting of multiple locations. Population activity was organized hierarchically, with neural distances between elements of a category being smaller than the distances between categories. At the single cell level, individual medial prefrontal cortex (mPFC) neurons were tuned to category and space independently, consistent with hierarchical population geometry emerging from random mixed selectivity, while neurons exhibited preferences for representing space or time across categories, consistent with functional specialization. These distinct selectivity rules across different levels of abstraction suggest that different aspects of prefrontal representations have disparate origins and may enable modularized computations across multiple behaviors and contexts.

## INTRODUCTION

Prefrontal cortex (PFC) is critically important for the organization and execution of behavior, including sensory integration, decision making, and cognitive control^1,2^. In particular, its functions are believed to be supported by networks of cells representing information that is currently relevant for the task at hand. The currently relevant information can include sensory or behavioral variables as well as information that requires abstraction from sensory information, such as behaviorally relevant categories and the rules of the task to be performed in the present context^3–5^ – all of which have been shown to be represented in prefrontal cortex.

Previous work has identified two major themes regarding how the PFC represents these variables. First, individual neurons in PFC and other areas have often been found to encode not a single variable, such as a particular stimulus or feature, but instead be selective for multiple variables, termed “mixed selectivity”^6^. Second, population vector activity has been found to exhibit structure corresponding to geometrical^7–9^ or hierarchical^10,11^ relationships among features or variables in the task. Such geometric structure, including the degree to which independent variables occupy factorized subspaces, has been suggested to govern downstream readout generalization^12^.

A key question is whether there exists any structure in terms of which variables each PFC neuron represents. For instance, in the case of mixed selectivity, is the selectivity for each variable randomly distributed across neurons? It has been found that neurons in orbitofrontal cortex and anterior lateral motor cortex can be grouped into clusters with similar response profiles, i.e. neurons that are selective for similar sets of variables^13–15^ (but see^16^ for different results in parietal cortex), and theoretical work has explored the benefits of such functional clusters as well as the conditions under which they might arise^17,18^. However, it remains unclear what kinds of functional preferences exist, how they relate to population level representations such as hierarchical structure, and whether the rules governing selectivity differ across levels of that hierarchy.

To address these questions we leveraged a spatial navigation task in which rats navigated between two behaviorally distinct location categories – starts and goals – each comprising three individual locations^10^ (Fig. 1A). We previously showed that medial prefrontal cortex (mPFC) population activity was organized hierarchically with respect to these categories: population vectors clustered by category, with starts grouped together and goals grouped together, while individual locations remained distinguishable within each category (Fig. 1B)^10^. This hierarchical population structure, combined with the known selectivity of individual PFC neurons for spatial locations^19,20^ and categories^3^, allows an investigation into how single cell selectivity for space and category gives rise to such population-level structure.

**Figure 1.**
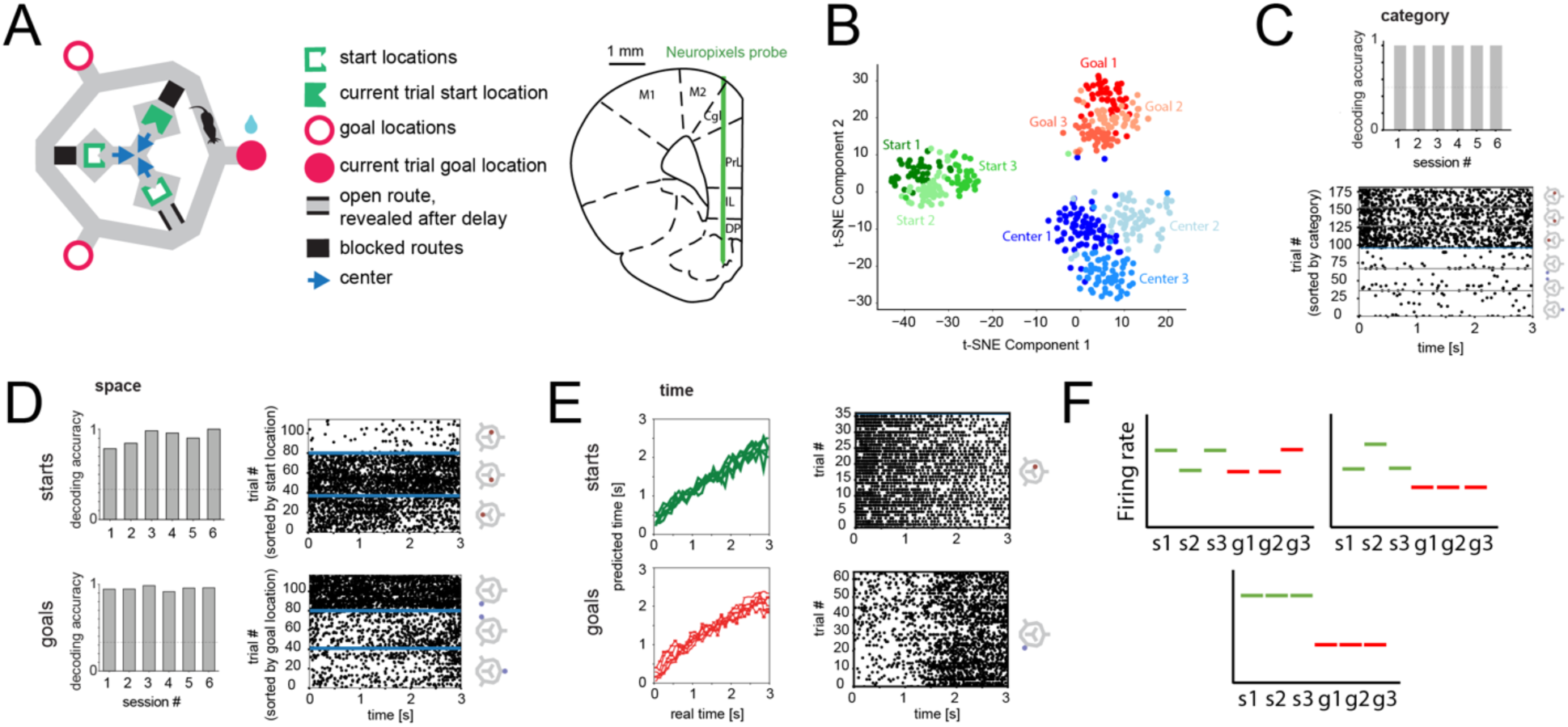
Hierarchical representation of population activity, with space and time represented in distinct location categories. **(A)** Navigation task layout (left) with two location categories (starts, green; goals, red) and recording location (right) Blue arrows indicate position and head direction of rat when returning to center via one of the routes. (**B)** Neural population activity of locations is hierarchically organized according to the meaning in the task, with clustering of starts, goals, and center locations, followed by division of those clusters into individual locations. **(C)** Start and goal categories can be decoded at the population level (top) (chance, dotted line). (**D)** Individual locations with start and goal categories can be decoded from neural population activity (left) (chance, dotted line). (**E)** Time can be decoded from population activity at starts and goals (left) (time bins 100 ms or, for two sessions, 106.7 ms, i.e. 30 bins for the 3 or 3.2 s delay period at starts). **(F)** Examples of hypothetical single cell firing patterns that may contribute to the observed spatial selectivity and/or hierarchical organization of categories (s: start, g: goal). Each raster plot in C-E shows an example neuron with the indicated selectivity across trials (with trials grouped as shown).

In addition to space and category, the PFC has been shown to encode time, for instance in temporal discrimination and reproduction tasks^21–23^ or during delays^24,25^. Time and space are fundamental variables present in all experiences and are critical for organizing and executing actions at the appropriate moments and places. Thus, we further investigated the distribution of selectivity for these key variables by analyzing how the selectivity of individual PFC neurons for locations and time were organized within and across behavioral categories.

Using simultaneous large-scale recordings from mPFC via chronically implanted Neuropixels probes^26^, we discovered that the rules governing selectivity differed across levels of the task hierarchy. We found that selectivity for specific spatial locations and for category were independently distributed across neurons, i.e. the hierarchical population geometry emerged from randomly mixed single cell selectivity. On the other hand, selectivity for spatial locations and time were not independently distributed across neurons. Instead, neurons were functionally specialized to represent spatial locations or time, as these neurons preferentially represented a given variable across category boundaries. Moreover, at the population level, the temporal code itself was stable across individual locations within a category, pointing to a generalized, location-invariant representation of elapsed time, and the spatial code exhibited additional task-related geometric structure. Thus, our results reveal multiple layers of structure in mPFC activity. These findings (i) provide new insight into the links between single cell and population-level coding, (ii) demonstrate distinct selectivity rules pointing to potential underlying anatomical connectivity patterns and/or other kinds of cell type specialization, and (iii) offer a framework for understanding how representations of different types of variables are structured. This organization could facilitate neural computations important for solving complex behavioral tasks under a variety of conditions.

## RESULTS

### Space, time, and category are represented in mPFC

In our delayed match-to-sample (multi-start/multi-goal/multi-route, i.e. “MSMGMR”) working memory task^10^, rats navigated from one of three start locations to one of three memorized goal locations using variable, unpredictable routes (Fig. 1A). During the sample phase of each trial, animals were visually guided to the randomly assigned current goal location and had to maintain that location in memory while it returned to the center of the maze and subsequently searched for one of three randomly assigned start locations. Once found, the animal had to nose-poke for 3 s (or 3.2 s in one animal) at this start location. During this time the animal was unaware which route would become available and thus could not plan its motor actions until the end of the delay period (Fig. 1A). At the end of the delay, a route was made available, and, during this test phase of the trial, the animal needed to navigate to the correct memorized goal location to obtain reward. In the next trial, the goal and start locations were again randomly assigned. Animals reached equally high levels of performance for all goal/start/route combinations (77.37% correct on average, chance: 1/3)^10^. Using chronically implanted Neuropixels probes, we sampled a large number of mPFC neurons (68 – 97 simultaneously recorded neurons per session after stability criteria had been applied, mean: 86.7, six recording sessions from 3 animals, 2 sessions per animal) during task performance.

We previously reported that neural activity at different locations clustered according to their meaning in the task^10^ (Fig. 1B) and hypothesized that such representations of categories are important for animals to understand and solve the behavioral task. The representations of these categories are hierarchically arranged, such that the distance between population vectors of the furthest elements within each category is smaller than the distance between categories (Fig. 1B). The goal and start locations are the most important locations in the task and the behavior at these locations was well-controlled; thus, we focused our analysis of neural selectivity for category, space and time on the delay periods at the three start locations and the periods at the three goal locations during the sample phase during which times the animals were stationary (see methods for details). Goal and start location categories could be perfectly decoded from population activity in all sessions (Fig. 1C, top) and single cells could be selective for category without spatially coding for individual locations within either category (Fig. 1C, bottom). Mean firing rate over all neurons did not differ significantly between delay and goal periods (mean difference: −0.2 Hz, range: −0.5 to +0.2 Hz across sessions; Wilcoxon signed-rank test, p = 0.22).

Spatial selectivity in mPFC has been described in open field settings and simple mazes^19,20,27,28^. In our maze, each of the six locations could be decoded from neural activity with high accuracy (Fig. 1D). The mean firing rate of an individual cell could differ for each individual location within one category, or one location could be different from (higher or lower than) two others (Fig. 1D, right, Fig. S1).

Neurons in PFC have also been shown to reflect elapsed time in a variety of circumstances^21,25,29,30^. In our task, at each of the start and goal locations, i.e. during the delay period at the start, and in the 3 s period after animals arrived at the goal, elapsed time could be decoded with a population decoder well above chance and single cells exhibited temporal modulation at start and goals (Fig. 1E).

Having established decodable neural representations of category, space, and time, we asked (i) what firing patterns at the single cell level underlie the hierarchical population level representation of starts and goals (e.g. are separate subsets of neurons selective for category per se and for locations within a category) (Fig. 1F) and (ii) whether selectivity for spatial location and/or time elapsed is maintained across category boundaries (e.g. is a neuron whose firing rate is modulated by the identity of the current start location more likely to be also spatially selective for goal locations, i.e. does PFC exhibit functional specialization).

### Distribution of individual cell selectivity for category, space, and time in mPFC

To assess how space and category are represented at the single neuron level, we defined cell selectivity for category in accordance with the hierarchical organisation observed at the population level: to be category-selective, a cell had to have a significantly different firing rate at starts versus goals, and the firing rate difference between the elements of each category had to be smaller than the difference between categories (see methods for details).

Out of 520 cells in total, the number of spatially selective cells at either starts or goals was 374 (71.9%), 271 (52.1%) were selective for category and also spatially selective (i.e. distinguished either between the three start locations or between the three goal locations), and 108 were exclusively selective for category (Fig. 2A, see also Fig. 1C and F and Fig. S2 for single session results). The number of cells selective for both space and category closely matched that expected by independent incidence (271 observed, 273 expected, p = 0.67), and similarly for exclusive category selectivity (108 observed, 106 expected, p=0.41). Moreover, category and spatial tuning effect sizes for individual cells revealed that the population code is made up of a range of different combinations of selectivity, including pure spatial and pure category selective cells. Cells could be strongly modulated by category but have strong or weak spatial modulation (Fig. 2B).

**Figure 2.**
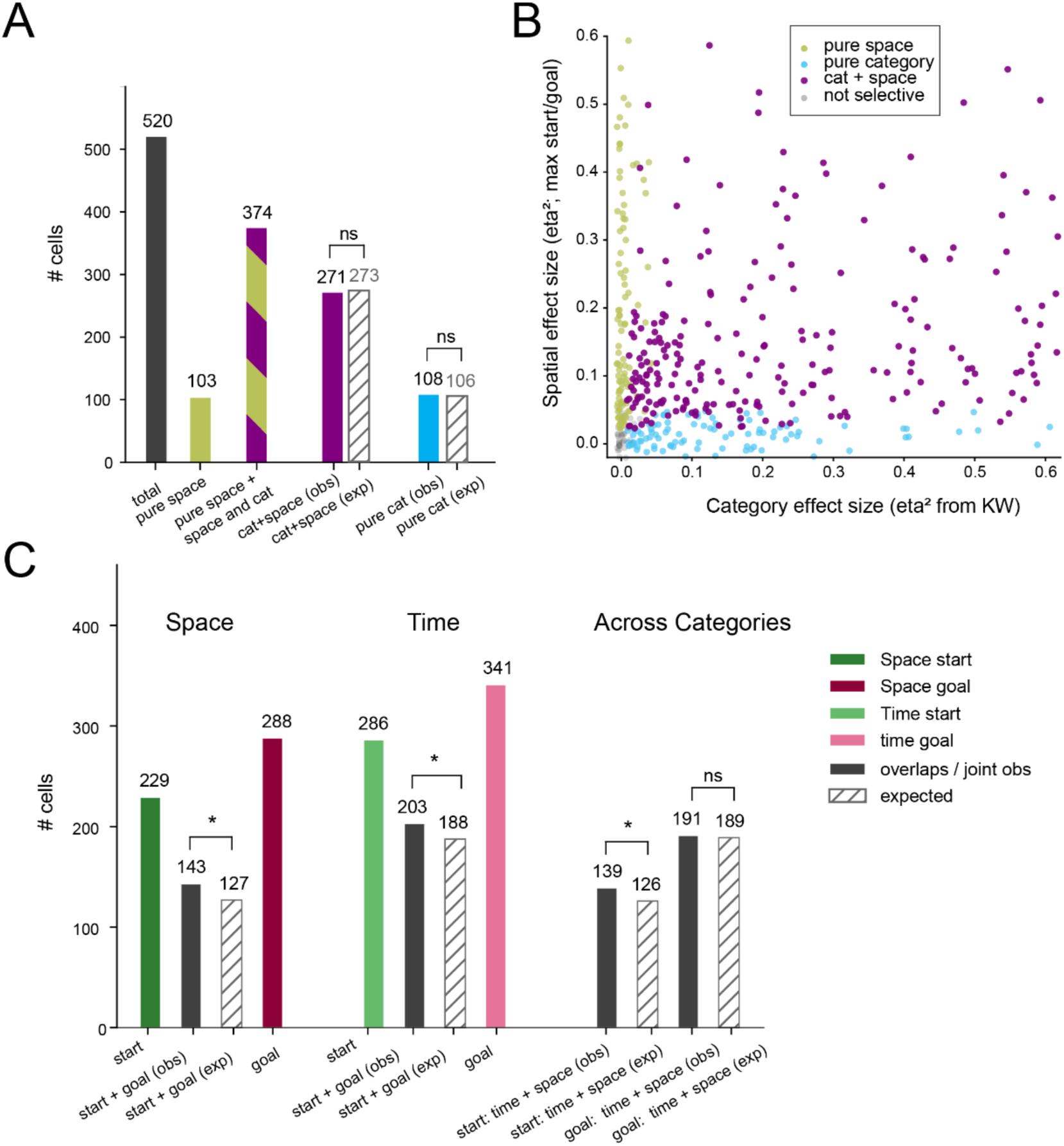
Distinct distributions of single cell selectivity for category and space compared to space and time across categories. **(A)** Number of cells selective for space only (‘pure space’), category only (‘pure cat’), and for various combinations. Number expected assuming selectivity for space and category are independent (hatched). **(B)** Effect sizes for category selectivity and spatial selectivity across cells. **(C)** Number of cells selective for space and time at starts and goals. Observed and expected (hatched) overlap assuming independence (N = 520 cells total). Kruskal-Wallis test was used to determine the number of significantly modulated cells. Fisher’s exact test was used to compare observed vs expected. *p<0.05

To investigate how individual neurons were modulated by space and time *across categories*, we compared each neuron’s spike count at the starts or goals for space, while for time elapsed spiking was binned into 3 time bins at each start or goal. Out of the 520 neurons, 229 (44.0%) were significantly modulated by space at starts and 288 (55.4%) at goals (Fig. 2C). The expected number modulated by space at both starts and goals if spatial selectivity at starts and goals were independent of one another would be 127 (24.4%). Instead, 143 cells (27.5%) (p = 0.0045) had both types of spatial selectivity, indicating a modest functional preference for encoding space in a subset of cells. Similarly, for cells modulate by time elapsed, we found 286 (55.0%) and 341 (65.6%) cells at starts and goals, respectively. The expected number of time-selective cells at both categories was 188 (36.2%), but we found a moderately larger number, 203 (39.0%) (p = 0.0053) (Fig. 2C). Thus, in contrast to independent selectivity for space and category described above, these results provide evidence that mPFC neurons have consistent functional selectivity that is maintained across category boundaries.

Are cells encoding space in one location category more or less likely to also encode time in that category? The number of cells encoding both space and time at starts was moderately higher than expected assuming independence: (139 (26.7%) observed, 126 (24.2%) expected (p = 0.0128). At goals, the observed and expected numbers were similar: 191 (36.7%) observed, 189 (36.3%) expected (p = 0.3801) (Fig. 2C). Thus, space and time selectivity were not exclusive; instead, many cells had mixed selectivity, i.e. were modulated by both space and time.

### Population decoding analysis of the interaction of category, space, and time selectivity

As we found significant biases in selectivity across categories but not rank (space versus category) at the single cell level, we then analyzed these properties of PFC coding in more detail at the level of neural populations.

To assess functional preferences or random mixed selectivity, we used logistic regression classifiers to probe how accurately category, spatial location, or elapsed time could be decoded from sets of neurons (Fig. 3A). To quantify the overlap between populations coding for space and category, we first ranked neurons by their impact on classifying space, i.e. we sorted them according to the highest absolute classifier weight from two spatial decoders, one trained to decode space at starts and, the other, space at goals. We then started with the one neuron with highest absolute weight and successively added neurons with decreasing absolute weight to build classifiers trained to decode category. Naturally, decoding improved with more neurons (Fig. 3B). For the top spatial decoding neurons, however, the ability to decode category was no better than a random choice of neurons, and the spatial and random curves were close throughout (Fig. 3B), consistent with our finding at the single cell level (Fig. 2A, B).

**Figure 3.**
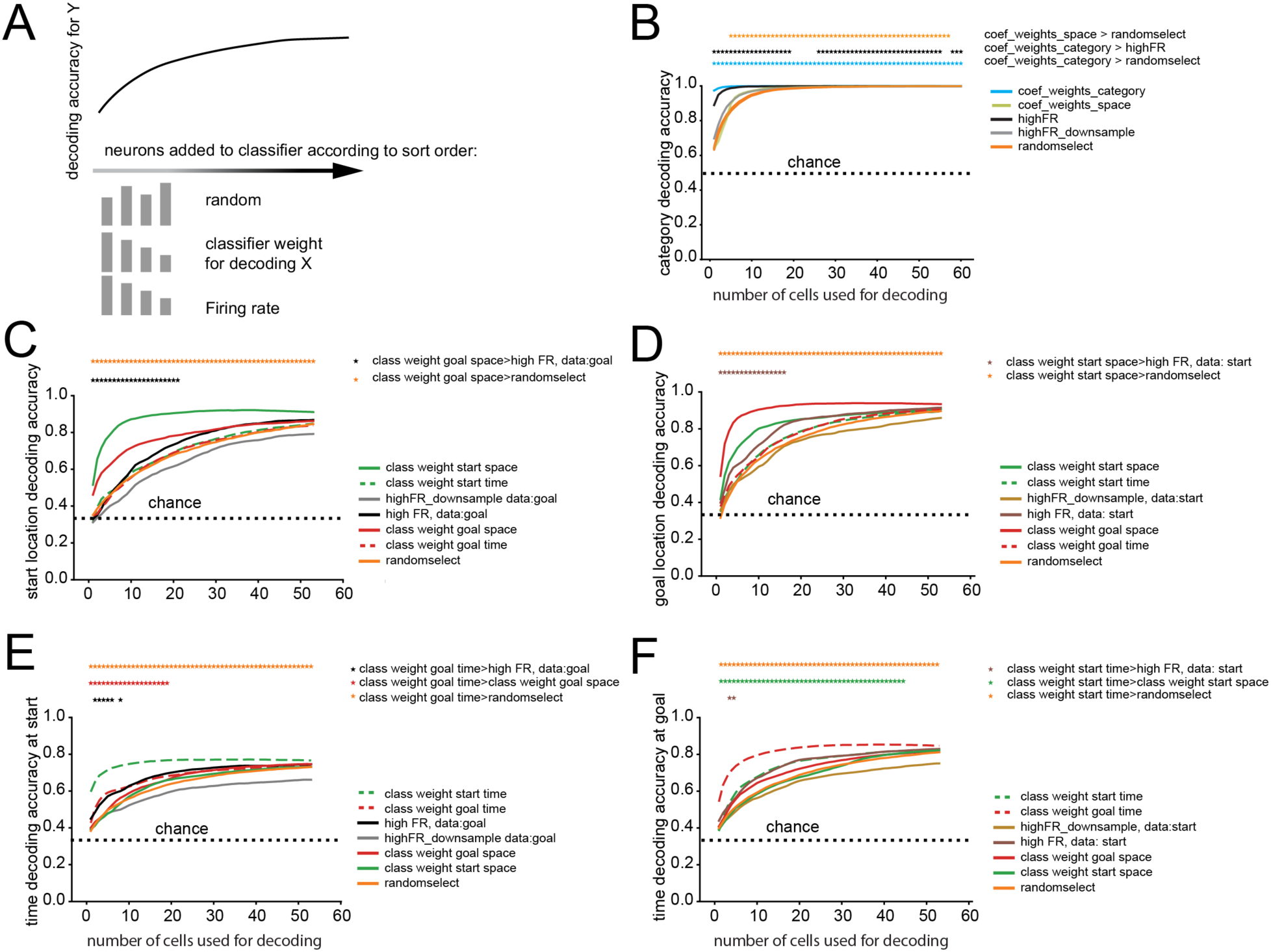
Population-level decoding analysis: functional preference of cells for space and time coding across categories and independence for space and category coding. **(A)** Schematic overview of analysis. Neurons are added successively according to different sorting rules to test their impact on logistic regression decoder performance. Decoding accuracy for category **(B)**, space (**(C)**, start, **(D)**, goal) and time **((E)**, at start, **(F)**, at goal) using increasing numbers of neurons. The order in which neurons were added to the respective decoder was based on classifier weights for space (solid green: start, solid red: goal, and, in **(B)**, yellow-green: both), time (dashed green: time at starts, dashed red: time at goals), category (blue), firing rate (at goal and at goal and start: black, at start: brown), or averaged downsampled firing rate (at goal and at goal and start: gray, at start: light brown). The orange line is the averaged decoding accuracy for 50 randomly chosen neuron orderings. For category decoding **(B)** and space decoding (**(C)** and **(D)**), the summed spike count during the full delay period at the start or a 3 sec period at the goal for each trial were used for classification. For time decoding (**(E)** and **(F)**), the analyzed time period at the starts and at the goals was divided into 3 time bins. Accuracy was determined as a binary decision. Stars above each plot denote multiple comparison-corrected significance (p<0.05).

We next applied an analogous approach to assess the overlap of neurons contributing to spatial decoding at starts (Fig. 3C) and goals (Fig. 3D). We sorted neurons according to the highest absolute classifier weight for any of the three goal (start) locations, then started with the one neuron with highest absolute weight and successively added neurons with decreasing classification importance to decode start (goal) location (Fig. 3C, D). Importantly, decoder performance for start locations improved more steeply when adding neurons ordered by goal decoder weights compared to randomly selected neurons (Fig. 3C), and vice versa (Fig. 3D). These findings demonstrate that an overlapping set of neurons is engaged in decoding both start and goal locations, consistent with single cell-level findings (Fig. 2C). To assess if firing rate alone could explain this finding, we compared these curves to the improvement of start (goal) decoder performance when adding neurons ranked by their firing rate (high to low, i.e. including higher firing rate neurons first). Firing rate did not account for the steep improvement in performance we observed for neurons selected by classifier weights, though higher firing rate neurons did outperform randomly selected neurons.

Similarly, to analyze functional preference for time across categories, we successively added neurons ordered by their weights for decoding time at goals (starts) to a classifier for decoding time at starts (goals) (Fig. 3E, F). Neurons added based on decoder weights improved decoding accuracy faster (i.e. with fewer neurons) than randomly selecting neurons, showing functional preference to decode time in a subset of cells. Selecting neurons based on firing rate (high to low) again improved decoder performance faster than randomly selecting neurons, suggesting higher firing rate cells have higher decoding power. Nevertheless, consecutively adding neurons based on time decoding in the other category was still significantly better than adding neurons based on high firing rate, though this was limited to decoders with low neuron numbers, i.e. the ‘top decoders’ of time in the other category were more informative than the highest firing rate cells.

We also assessed individual neurons’ decoding potential across space and time (i.e. selecting neurons based on classifier weights from a space decoder to decode time and vice versa, Fig. 3C-F). The improvement in decoding accuracy when adding neurons ranked by classifier weights of the respective other variable was no different than with randomly ranked neurons. This suggests that preference to decode space and time overlapped no more than expected by chance, but also no less – i.e. preference for space or time was not exclusive.

As described above, ranking neurons by firing rate generally yielded steeper improvement in decoding accuracy than randomly ordered neurons, thus high firing rate was a common predictor of decoding ability for category, space, and time. We tested if high firing rate cells tended to be important because of a potentially higher signal-to-noise ratio, or because these cells were more potent predictors per se. We reduced the number of spikes of these neurons by downsampling the number of spikes to match the firing rates of randomly drawn neurons (see methods for details). The resulting decoding accuracy curve was lower than or similar to that of randomly ordered neurons, suggesting that firing rate itself was responsible for this effect and not another property of these cells (Fig. 3 B-F).

Taken altogether, these results reveal an organization in mPFC where (i) neurons maintain their functional preference for space and time across category boundaries, while (ii) a different picture emerges for decoding category, where subsets of neurons independently code for space and category.

### The time code is location invariant but category-dependent

We next asked the following question: since cells are functionally specialized, is the firing rate modulation (i.e. the code itself) transferrable or related between locations and/or categories? For example, is the time code at start location 1 related to that of start location 2 or even the time code at the goal locations? To answer this question, we first visualized the relationship between neural activity at the six different locations as time elapsed using principal component analysis (PCA) of the average population vectors at these locations (Fig. 4A, Fig. S3). The variance explained by the first principal component was on average 5.5 times higher (range: 3.1 to 7.5, Fig. 4A) than the second and appeared to largely represent the overall difference between neural activity at start versus goal locations, consistent with previous analysis using t-SNE (Fig. 1B) and in line with a hierarchical representation, where the difference between categories is larger than that between elements within one category. Thus, we performed PCA separately for the goal and start locations to visualize the relationship between spatial and temporal components (Fig. 4B and Fig. S3) within a location category. Here, the temporal trajectories were separate for each location and had a common shape, unfolding in parallel for the separate locations. Location and time representations were primarily represented in principal component 1 and 2, respectively, and thus were orthogonal to one another. This points to a common, generalized representation of time at different start and goal locations.

**Figure 4.**
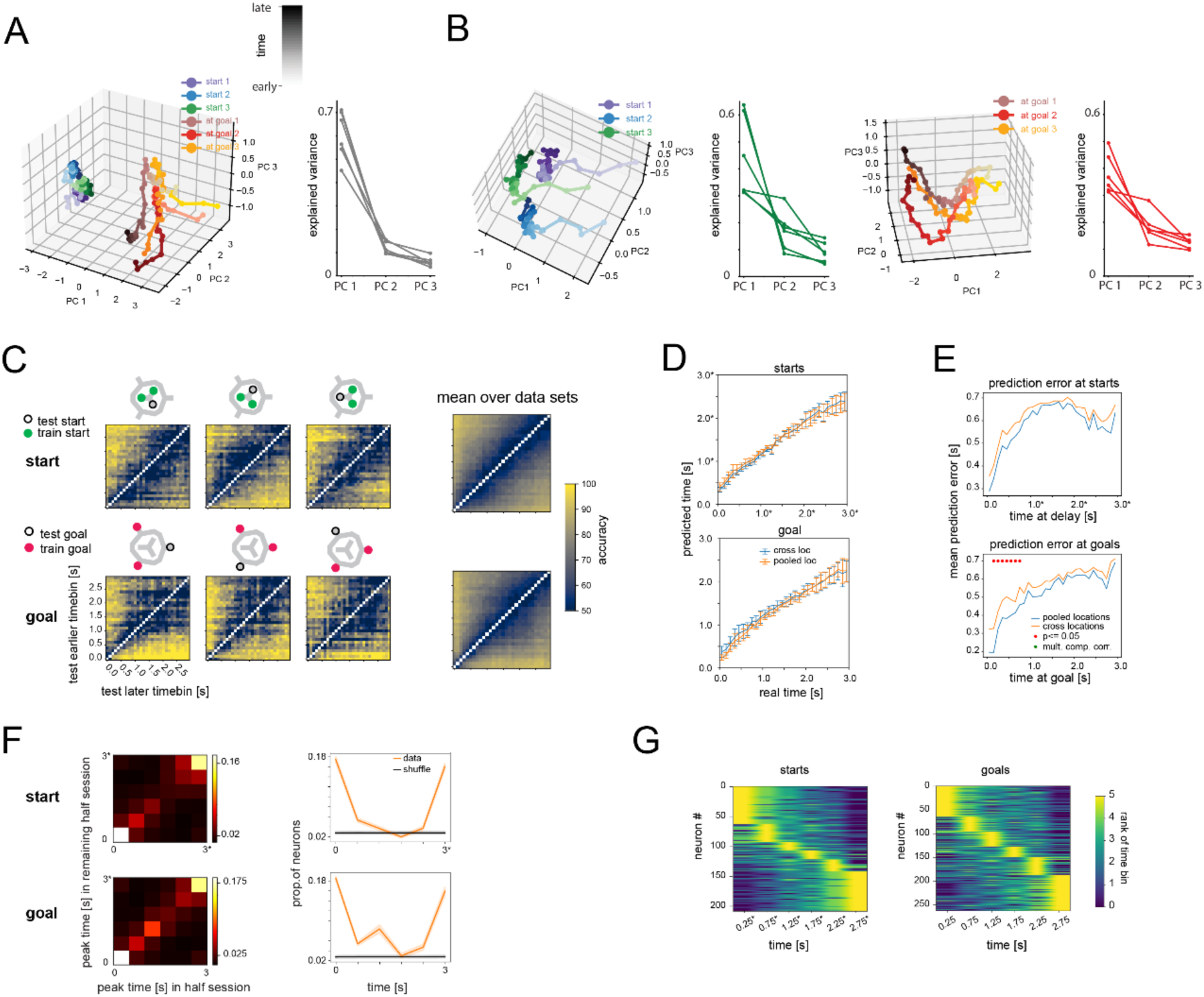
A generalized time code in prefrontal cortex. **(A)** Principal component analysis (PCA) of mPFC population activity for one session (averaged over trials) at all six locations (starts and goals) together. Time bins: 100 ms. Right: variance explained by the first three PCs for all sessions. (**B)** PCA and explained variance for start (left) and at goal (right) locations separately. Time evolves in parallel for different locations and primarily along one principal component. Color shades from light to dark correspond to early to late time points at the starts/goals (as indicated in grayscale in A, top). (**C)** Binary classification of which of two time bins neural data comes from for all possible time differences for starts (top) and goals (bottom). Maze symbols indicate for each row the two locations used for training and the test locations. Results above (below) the diagonal correspond to prediction accuracy of the later (earlier) of the two time bins. (**D)** Time decoding using all time bins, for training and testing across different locations (blue) or train trials and test trial were drawn from pooled locations (orange). Error bars are standard deviation (std) across sessions. (**E)** Prediction error (of decoding using all time bins). (**F)** Consistent activity peaks are primarily at the beginning and at the end of the analyzed time periods: left, comparison of activity peak when dividing dataset in halves; right, proportion of cells with a consistent activity peak at indicated time points (diagonal of matrices on the left, see text for details). (**G)** Cells ramp up or down at starts and goals. Color indicates for each cell the order (rank) of mean firing in each time bin (only cells with consistent peaks are shown, cells are ordered by the peak firing time bin). Displays show pooled results across all data sets for **D** and **E**. Time bins are 100 ms (and *106.7 ms for two of six sessions at starts) for A – E and 500 ms (and 533.3 ms for two of six sessions) for **F** and **G**.

We tested this hypothesis of a common time representation directly by training neural population decoders to decode time using any combination of two locations within a category and testing their performance on the respective third location. For this we used binary logistic regression decoders that were trained on the neural data of any combination of two time bins (Fig. 4C). Decoder accuracy of which of the two time bins the data was from was close to perfect for time bins temporally furthest apart and lower for time bins closer in time. We further verified these findings by using all time bins and training and testing across locations, and assessing decoding precision (Fig 4D, E).

Taken together, not only are prefrontal cortex single neurons specialized for time, but there is a common population code that reflects elapsed time across different locations within each category (goal or start).

Given that the time code is resistant to interference by the spatial location code, what firing rate modulation might differentiate time points at the single cell level? One possibility to signal elapsed time that may lead to the representation at the population level described above has been described in parietal cortex and other brain areas: tiling of the delay period by neurons with firing peaks at different time points^31^. We used a previously described method^32^ to determine if cells had consistent firing peaks (Fig. 4F). Specifically, we repeatedly randomly divided each data set into halves and determined the peak of firing in the first half and compared it to the timing of the peak found in the second half. Most cells with consistent firing peaks peaked at the beginning or at the end of the considered time period at starts and goals. These cells tend to increase or decrease their firing gradually (Fig. 4G), consistent with a time code that is primarily generated by cells’ firing ramping up or down instead of sequential firing.

We then tested whether the time code was transferrable across goal and start categories. Results differed for different sessions. In one case the time codes for goals and starts overlapped partially, in two cases they were partially reversed, and in the remaining three sessions we saw no evidence for cross-category time decoding (Fig. S4). The time codes are thus generally not consistent across start and goal locations even though the contributing cells are overlapping (Fig. 2C, Fig. 3E,F).

### Geometric representation of spatial locations

As with time elapsed, we next considered whether there might be additional structure for the spatial code. We explored potential structural relationships across categories using a decoder trained to predict spatial locations from one category (e.g. start locations) and testing it on spatial locations from the other category (e.g. goal locations). This should yield random predictions if spatial codes for start and goal locations are unrelated. Instead, we found an unexpected reflection of the spatial layout of the task: the goal location opposite the start location in the training set was predicted far below chance in all sessions (Fig. 5A; see Fig. S5A,B for the reverse analysis, i.e. training on goal data and testing on start data, and for single session results). Thus, the classifier axes dividing the start locations in neural activity space divide the respective goal locations into opposite and adjacent goals. This points toward more similar population vectors between goal and start locations that are not located opposite of one another. We tested this directly by correlating the population vectors at start and goal locations with one another (Fig. 5B, Fig. S5C). Indeed, correlations (after standardization) were generally negative if the start and goal locations were opposite of one another, and positive or uncorrelated if they were adjacent. We found similar results when we correlated the weights from classifiers trained separately on start and goal locations (Fig. S5D). These results demonstrate an unexpected geometric relationship across location codes for behaviorally meaningful categories that is defined by their relative location in physical space. One possible explanation for the relationship between opposite and adjacent start and goal locations could be their direct physical distance in space. This distance was larger between a start and its opposite goal location than between a start and its two adjacent goal locations. However, there were no direct connections between these locations in our elevated maze and the distance an animal traversed from a start to the opposite goal could also be shorter than to an adjacent goal depending on the available bridge in a given trial. Other possibilities could be similar visual inputs or the influence of head-direction tuning. The animal’s head direction is 60 degrees offset for the goals adjacent to a start and 180 degrees for a start’s opposite goal location. Thus, cells with broad, unidirectional tuning could contribute to our finding that start and goal location codes are related.

**Figure 5.**
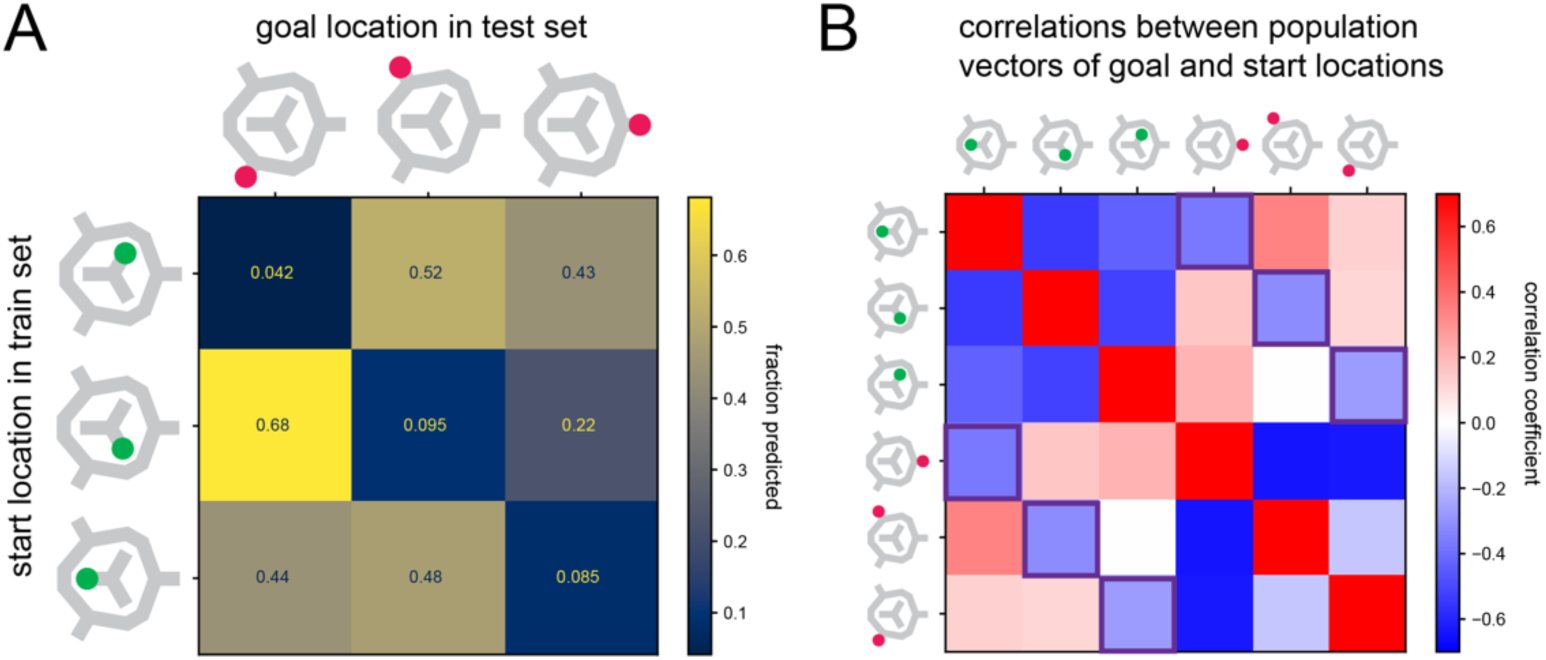
Spatial codes across categories are structured. **(A)** Predicting goal location with a classifier trained on start locations. Opposite locations were rarely predicted. Mean over sessions. (**B)** Correlation coefficients of population vectors of start and goal locations (the goal location and start location population vectors of all trials were standardized separately before correlation to reveal any structure beyond the high similarity among start and goal location population vectors). Mean across all 6 sessions. The squares with dark edges denote values corresponding to opposite start and goal locations.

Taken together, for spatial codes we found that, in addition to functional specialization, there was a structured relationship across location categories.

### Space, time, and category selective cells are distributed across all subareas of mPFC

Having identified cells with distinct functional preferences and additional structure within and across categories, as well as cells that code for category itself, the question arises whether cells with any of these functional features might be organized anatomically. Thus, we tested whether neurons from subregions of the mPFC (which were all recorded simultaneously) were differentially organized depending on how strongly they were involved in decoding category, or, spatial or temporal variables. We compared the distribution of classifier weights from space, time, or category classifiers and tested if there was any bias that could be explained by the recording location (Fig. S6, S7). We did not find any consistent differences in anterior cingulate cortex, prelimbic or infralimbic cortex, or dorsal peduncular cortex (see methods for details). However, it should be noted that the precision of our anatomical evaluation of the recording locations was limited, and results for other methods and variables can lead to different results^33^.

## DISCUSSION

The central role of frontal cortices in goal-directed, cognitively demanding tasks is well-accepted: in working memory tasks, prefrontal cortex is required for all phases of the task^34^. A multitude of neural correlates of task-related variables at the population and single cell level that presumably enable prefrontal cortex function have been described. A fundamental question is how single cell selectivity and population level representations are linked and whether single-cell selectivity for particular variables is governed by specific rules. Here, we addressed these questions by focusing on neural correlates of space and time and their relationship to a third, higher-order variable, location category, which served as the behavioral category in our task and was also represented at the population and single-cell level.

Categorization is a key aspect of prefrontal coding^3,5^. In our task, the behavioral distinction of the two categories, starts and goals, was implemented as nested clusters in neural activity space: the distance between categories was larger than the distance between individual locations within each category. Thus, while a representation where within-category and between-category distances are equal could have still supported categorical discrimination, we instead found that the neural structure mirrors the hierarchical nature of the task variables. This suggests that animals did not merely learn to distinguish starts from goals, but internalized the nested organization of the task, i.e. that individual locations are instances of a broader category rather than simply distinct places. At the single cell level, category was represented both by cells that also showed spatial selectivity and by pure category cells.

With these two spatial categories, we could also probe the same neurons across category boundaries. This internal comparison allowed us to discover that subsets of neurons, while often displaying selectivity for more than one variable (mixed selectivity), preferentially encoded space or time elapsed across behavioral categories. Thus, even though the behavioral category at starts or goals was different – a difference that was also clearly reflected in neural activity and represented at the population and single neuron level – selectivity for a particular variable was not independent of the two categories as might be expected if selectivity were dynamically and randomly assigned in each category. A similar approach has been taken in an economic decision-making task in monkeys^35^. Here, animals were presented with two different sets of choices, and orbitofrontal cortex (OFC) neuron selectivity for economic decision variables was compared across those sets of choices. Neurons maintained their specific selectivity across choice sets and thus formed a stable framework that can adapt to different contexts by ‘remapping’ their selectivity to the currently relevant choice set. These findings reveal functional specialization, i.e. a neuron’s selectivity is not randomly assigned or determined by ongoing activity, but instead is consistent across different contexts and thus appears to be a property of the cell. A study in medial entorhinal cortex (MEC) investigated selectivity for space and time of individual neurons across visually different environments^36^ and, in line with our results from mPFC, MEC space and time cells preferentially encoded the same variable across different contexts. Furthermore, in MEC, space and time cells were organized in anatomical subclusters and associated with different behavioral states (movement and immobility). Beyond consistency of selectivity across conditions, functional specialization has also been explored as a question of network organization, i.e. whether neurons are organized into functional subclusters that have similar or overlapping selectivity profiles. Using unbiased clustering approaches, the organization of single neurons into functional groups has been revealed in rodent OFC^13,14^. Hirokawa et al.^13^ further linked a group of functionally distinct neurons to a specific projection target, suggesting that functional separation might be accompanied by distinct connectivity. In mouse anterior lateral motor cortex (ALM), functional subpopulations were also found and linked to the varying strength of input from other brain areas^15^.

Unlike selectivity for space and time, selectivity for category itself was distributed across an unbiased, effectively random subset of cells in our task. Thus, even though the categories here have a spatial dimension, there is a dissociated from which cells code for specific spatial locations. This suggests that the principle underlying selectivity for more abstract variables may be distinct from that of sensory or internal variables, and that functional preference does not cross ranks in a hierarchical representation.

At the population level, we also found that the code for elapsed time in prefrontal cortex is flexibly transferrable to different locations within location categories (i.e. between start locations or between goal locations). Thus, a neuron’s firing rate modulation as time elapses (in most cases ramping up or down, Fig. 4F, G) is not influenced by the current location of the animal, if those locations are of the same type, i.e. within one category. This is in line with a finding in parietal cortex where choice and modality were coded independently of one another^16^. Thus, while prefrontal cortex neurons are often selective for multiple variables, behaviorally important variables can be independent of one another.

While we found that the time code is transferable within one location category, a largely different code is used for the respective other category, even though there is an overlapping set of time-selective neurons for both categories. Note that the delay period (at starts) is a fixed time interval, while goal timing is tied to the onset of a behavioral event (i.e. arrival at the goal). Furthermore, rats received a reward at the goal, but not during the delay period. Therefore, other factors in addition to elapsed time likely impacted the neural dynamics differentially across these categories. Of note, animals also receive a small reward when returning to center, but the representation of this rewarded time period did not cluster with the goal (Fig. 1B); thus, reward itself or overall average firing rate, is not responsible for clustering.

Interestingly, the spatial representations across the two location categories preserved the geometric structure of the maze: start location codes separated the opposite goal from the adjacent goal locations, mirroring the physical distances between locations. This is consistent with a broader principle in which prefrontal neural geometry reflects the relational structure of the task, as demonstrated for abstract task variables in PFC and hippocampus^7^, goal locations in OFC^8^, and ordinal sequences in frontal cortex^9^.

Structural organization into functional subclusters, which often implies functional specialization, has been proposed to be computationally beneficial or even required for cognitive flexibility, as for example when learning multiple tasks and transferring learned representations from one task to another^17^. In artificial networks trained to perform common neuroscience tasks, those that required flexible input-output mappings displayed non-random population structure^18^.

Future work should examine further characteristics of the functional organization of mPFC, such as the identification of subsets of cells that are defined by projection targets or molecular markers^13,37–39^. Another area of investigation is how representations of task-related variables – abstract, sensory, and internal ones - and functional specialization emerge during learning at the population and single cell level, how single activity is coordinated during task execution trial by trial, or whether there are factors that predetermine such functional organization. The distinct organization schemes linking single cell selectivity and population level representations that we found could modularize functional elements of behavior and may be a hallmark of brain areas involved in higher-order cognitive functions.

## MATERIALS and METHODS

### Experimental procedures

Details on experimental procedures including surgery, behavior, and histology have been described previously^10^. The data used for the current study partially overlaps with our previous work where we reported a different set of findings on separate scientific questions. All procedures were conducted in accordance with the Janelia Research Campus Institutional Animal Care and Use Committee. Briefly, the data presented here was collected from three male Long-Evans rats. From each animal, 2 sessions from two days that were 3 to 4 days apart were included (the first but not second session from each animal was analyzed in^10^). Single-shank Neuropixels 1.0 probes^26^ (https://www.neuropixels.org) were implanted chronically and fixed in place (i.e. not moved in between recording sessions). In each animal, one such Neuropixels probe was implanted in the right hemisphere at 3.24 mm anterior-posterior and 0.6 mm medio-lateral with respect to bregma, at a depth of 6 or 6.2 mm below the brain surface. The recording sites spanned anterior cingulate cortex, prelimibic and infralimbic cortex and, in 2 animals, dorsal peduncular cortex. The probe location was confirmed with standard histological procedures. Recordings were conducted once animals were fully trained in the multi-start/multi-goal/multi-route (MSMGMR) behavioral task. This match-to-sample spatial working memory task was conducted on an elevated maze and consisted of a sample phase and a test phase. In the sample phase, the animal navigated from the center (Fig. 1A) to one of the three goal locations (pseudorandomly (i.e. balanced) chosen in each trial and visually marked to guide the animal via the available bridge (out of three total bridges, which were movable elements in the elevated maze that connected the outer ring and goals with the center; the available bridge in the sample phase was chosen pseudorandomly). Once the goal was reached, a small liquid food reward (Ensure Plus) was automatically delivered. The delivery was triggered by the animal breaking an infrared beam at the reward pod. Next, the animal returned to the center, received another small reward, then had to search for the one location of the three start locations (Fig. 1A, green rectangular shapes) that was pseudorandomly chosen in each trial. Each of the start locations was equipped with a nose poke port, and the correct nose poke (i.e. the nose poke at the start chosen for that trial) was indicated by a tone emitted from the port while being poked. The nose poke duration (‘delay period’) was 3 s in two animals and 3.2 s in the third animal. During this period, all bridges were lowered, i.e. the animal had no access to the outer ring of the maze. Which bridge would eventually be accessible was not known to the animal and thus the animal could not plan or prepare the route it would have to take during the delay period. After the delay period at one of the start locations, a pseudorandomly chosen bridge became available and the test phase began. In the test phase the rat had to navigate, without any external cues, to the goal location that was rewarded in the sample phase. If the animal chose the correct goal location, it received a reward, then it returned to the center and the sample phase of the next trial began.

Neural data was recorded and stored using SpikeGLX software (http://billkarsh.github.io/SpikeGLX/). The spike data was first automatically presorted and then manually curated using the JRCLUST software package^40^. We isolated on average 142.8 neurons per session (range: 98 - 182) across prefrontal cortex subareas. Only cells that were stable over the recording session, selected based on previously described methods^10^, were used for the analyses shown in this study. After also excluding putative interneurons, the number of neurons analyzed was on average 86.7 neurons per session (range: 68 - 97), totaling 520 neurons over the 6 sessions. The core analyses were repeated and the results confirmed when including all clusters isolated during spike sorting.

### Data Analysis

All analyses were conducted using custom-written code in Python or MATLAB. The analyses presented here were focused on two defined periods during task execution. First, the delay period at any one of the start locations (3 or 3.2 s in duration) and, second, the period after the animal had arrived at any one of the three goal locations. For analyses where the spike data were transformed into a spike rate by splitting the time period into time bins and counting the number of spikes that fell into each bin, the binwidth for all analyses involving the delay period was adjusted to the length of the delay period, so that for all sessions the same number of bins was used. The time period used for analysis at the goal was 3 s, starting from the time point the infrared beam at the reward pod was broken. The reward was triggered by the animal breaking the infrared beam at the reward pod. The animal was free to leave at any time but generally remained at the reward site to consume the reward for at least 3 s. As this also matched the delay period duration, we chose to analyze neural activity in this 3 s time window at the goals. To reduce variability in the time period at the goal, we only used data from the sample phase but have repeated all major analyses using data from the test period. (We had previously shown that the activity at any one goal location during sample and test phases is distinguishable but overlapping and distinct from that at any other goal location during test and sample phases^10^.) For the delay period at the start locations, we used trials where in the following test phase the animal made a correct or incorrect choice. For the goal periods we used only trials where the animal chose the correct, visually indicated goal location and disregarded the outcome of the following test phase.

#### Decoding space, time and category

For all of the space, time or category decoding analyses we used multinominal or binary logistic regression classifiers with L2 penalty. Results were comparable using other methods, such as support vector machines (SVM) or, for time, using linear regression with continuous response variables. For time decoding, we used a variable number of time bins, as indicated in the Results or figure legends.

#### Selectivity

To test if cells were selective for space, time and/or category, we used the Kruskal-Wallis test on spike counts of the time period during the delay at start locations (3 or 3.2 sec) or at the goal (3 sec). For category and space, the full duration of these periods was used. To assess selectivity for time we divided the time at starts and goals in three equally sized bins. For category selectivity we employed another criterion to ensure that strong spatial selectivity at one location would not be mistaken for category selectivity: For each neuron, we computed mean firing rates separately for the delay and goal categories (spike counts per trial divided by that epoch’s duration). Within each epoch, we computed the mean firing rate for each spatial label (1–3) and the average within-category “element distance” as the mean absolute difference between these label means (*d_delay_* and *d_goal_*). We then computed the between-category distance as *|µ_delay_ − µ_goal_|,* where *µ* is the mean firing rate pooled across labels. A neuron passed the category criterion if *|µ_delay_ − µ_goal_| > 0.5 × (d_delay_ + d_goal_).* Thus, neurons were considered category-selective only if the separation between delay and goal firing rates was sufficiently large relative to the distance between elements within each category. This criterion ensured that category selectivity exceeded variability attributable to spatial tuning.

For all Kruskal–Wallis tests, we quantified effect size using eta-squared (η²) derived from the H statistic: *η²= (H - k + 1)/(n - k)*, where *H* is the Kruskal–Wallis test statistic, *k* is the number of groups, and *n* is the total number of trials across groups. For spatial selectivity, η² was computed separately for delay and goal categories, and the larger of the two values was used for visualization in Fig 2B. For category selectivity, η² was computed from the two-group (delay vs goal) Kruskal–Wallis test. Effect sizes were used in scatter plots to quantify the magnitude of spatial and category tuning independent of statistical thresholding. To assess if the overlap in distributions was different from that expected from independent distributions we used the Fisher’s exact test.

To analyze details of spatial selectivity (Fig. S1), we first determined for each data set the cells that had different firing rates at the three different locations within a category using the Kruskal-Wallis test. We then used pairwise post hoc comparisons (Conover’s test) to determine which pairs were different and compared the firing rates of those that were different.

#### Functional specialization – decoder analysis

To assess whether neurons were functionally specialized at the population level, we compared the accuracy of logistic regression classifiers (for category and space or time, at starts or at goals) while successively adding neurons sorted according to four different methods (Fig. 3):

- Classifier weights: As a measure for the importance of a particular cell to decode a particular variable, we used the highest absolute classifier weight for any of the classes in a classifier trained to decode a particular variable. Cells were then sorted from highest weight to lowest weight and added in this order to compare the impact on accuracy compared to another order. Five different cell orders were determined in this way, corresponding to the two variables we analyzed (time and space) and the two location categories (starts and goals) and category itself.
- Random selection: Here, cells were ordered randomly (without replacement).
- Firing rate: For each location category (starts and goals), the summed spike count at all starts or at all goals was used to order cells from highest firing rate to lowest firing rate. For category decoding the sum of starts and goals was used and then ordered from high to low.
- Downsampled firing rate: To match the firing rate of high firing rate neurons to that of randomly selected neurons, we first ordered neurons according to their summed spike count at goals or at starts, respectively, as above. Next, we generated a randomly ordered list of the same neurons. We then matched the firing rate of each cell in the rate-ordered list to that of the cell in the randomly generated list at the same position. For each pair of neurons at the same rank position, spikes were randomly removed from the higher-firing neuron until its spike count matched that of its randomly assigned counterpart. If the randomly paired neuron had an equal or higher spike count, no spikes were removed.

For the category decoding, the principle was similar but the order was determined from the summed count in both categories, while the downsampling would still be performed within a category.

To achieve reliable estimates of the resulting accuracy curves, we used 80% of the neurons available in each session, performed the different sorting procedures with these selected neurons, determined how accuracy improved with an increased number of neurons, then repeated this procedure 50 times and pooled the results from all sessions. We then tested separately each feature size (i.e. the number of neurons included in the classifier) and compared sets of different orders of neurons (e.g. weights for start decoding and random selection) using the Mann-Whitney-Test, the results of which were then corrected for multiple comparisons using the Holm-Sidak method. For Fig. 3, the results from the 50 runs and all six sessions were averaged.

#### Principal component analysis and location invariance of the time code

For the principal component analysis in Fig. 4 and Fig. S3, the data at the starts and at the goals were binned into 30 equally sized time bins, averaged over trials for each location, and centered.

To test if the time code was transferrable to different locations (Fig. 4C), we trained a decoder on two locations of one location category (starts or goals) and tested its performance on the third location of that category. Each combination of two time bins was trained and tested separately. For example, a classifier was trained on goal locations 1 and 2, time bin 1 and time bin 10 (30 time bins of size 100 ms, or 106.7 ms for the 2 sessions during the delay where the delay period was 3.2 s for one animal), and then tested on time bins 1 and 10 on data from goal location 3. The upper and lower half of the matrix in Fig. 4C are the prediction accuracies for the earlier or later of the two time bins, respectively. The diagonal would not be a valid comparison (one class only) and is thus not filled. All possible combinations of two time bins for all possible combinations of two training locations and one test location were analyzed in this way.

When we determined prediction accuracy over all time bins (Fig. 1E, Fig. 4D) we used three approaches. First, only data from one start or one goal location was used (Fig. 1E). Second, data from all locations of one category were pooled (Fig. 4D). In these two cases we trained on all trials, except for one which was used for cross-validated testing. The procedure was repeated for all test trials. Third, we trained using data from two locations and tested on the third (Fig. 4C, D). We repeated this procedure for all three possible combinations of two training locations and one test location for each category. Because here the test set was separate from the train set by design, we did not perform leave-one-trial-out testing but instead evaluated all test trials on the same trained model. For all approaches the mean predictions were obtained by taking the average across the predictions across all trials for each time bin. To assess differences in mean prediction error (Fig. 4E) we calculated the error (the absolute difference between predicted and true time) for all trials from all sessions and tested each time bin separately using the two-sample Kolmogorov-Smirnov test. P-values were based on two-sided tests and corrected for multiple comparisons using the Holm-Sidak method.

For time predictions across categories (Fig. S4), we trained a logistic regression classifier on all trials from one location category, i.e. all three start locations, and tested on all locations of the other category. We split the time period at the three start locations and at the three goal locations into three time bins. All data were standardized together, i.e. all time bins from both location categories were concatenated, the resulting mean was subtracted, and each time was divided by the standard deviation. We also tested different forms of standardization: standardizing data from each category separately or determining the mean and standard deviation from one category and applying the parameters to the other category. The results varied slightly for single sessions across these methods, but none of these standardization methods reliably facilitated time decoding across category boundaries for all sessions.

#### Time code – sequential activity versus ramping

To test if the underlying neural code for time was based on sequential firing, we employed a modified version of a previously described method^32^. For each session, the data at the goal or at the start was binned into six time bins and a random half of all trials was averaged (Fig. 4F). For each cell the time bin with the highest firing rate was found. Next, the other half of trials was averaged and again the time bin with the peak firing rate was determined for each cell. This procedure was repeated 1000 times and the proportion of neurons whose peak firing fell into the same time bin for both splits was determined. If there was more than one time bin with the same firing rate, one of them was chosen at random. As a control, we shuffled the time bins in all trials (1000 times) and performed the same procedure (Fig. 4F). As the majority of cells with consistent firing peaks were active at the beginning or the end of the analyzed period, we tested whether cells tended to ramp up or down, i.e. gradually changed the firing rate, which could serve as the basis for the observed time code. We restricted this analysis to cells which fired more consistently than 95% of the shuffled controls. For each cell we then determined the rank with regards to firing rate of each time bin and sorted them according to the time bin with the highest rank (Fig. 4G).

#### Cross-location category spatial decoding

For the cross-location category spatial decoding (Fig. 5, Fig. S5) we trained a logistic regression classifier on all trials from one location category (starts or goals). We used the trained classifier to predict locations in the respective other category and found that the predictions were not randomly distributed but biased towards predicting the respective adjacent location. Training a classifier on goal or start locations yielded three weight vectors for both categories, one for each class, with a length equal to the number of neurons recorded in that session. We calculated Pearson’s correlation coefficient for the six resulting weight vectors to examine the relationship between start and goal location codes (Fig. S5C). For the correlations of population vectors (Fig. 5 and Fig. S5C), we first standardized the data at goals and starts separately by subtracting the mean and dividing by the standard deviation. We then took the mean at each of the six locations resulting in 6 vectors and calculated Pearson’s correlation coefficient.

#### Subareas of prefrontal cortex

To determine whether any of the mPFC subareas we recorded from were more or less involved in coding for time, space or category, we determined the approximate recording location for each cell with reference to the surface of the brain (Fig. S6, 7). We trained classifiers to decode time or space at the start and goal locations or distinguish between starts and goals (category) (using 3 time bins for time and one time bin for space and category). The classifier training resulted in weight vectors for each of the classes (locations one to three, time points one to three, category one and two). The maximum absolute weight across classes for each cell was taken as a measure for its ability to distinguish between the classes for this variable. For visualization in Fig. S6 and 7, we divided the maximum absolute weight by the sum of all weights of all cells in that session. For the subarea comparison, we again used the maximum absolute weight across classes for each cell and pooled the results for each session for start and goal locations, resulting in two values for each cell for time and two values for space. The cells were split into groups corresponding to the mPFC subareas. For all variables (space, time, category) we used the Kruskal-Wallis H-test to determine whether the medians of the groups were equal. In sessions where they were not equal, we further identified the groups that were different using post hoc pairwise tests for multiple comparison (Dunn’s test). We found no significant differences that were consistent across sessions and areas.

## Acknowledgements

This work was funded by the Howard Hughes Medical Institute. We thank S. Erwin and R. Gattoni for help with animal training, T. Harris, J. Colonell, B. Karsh and W. Sun for help and advice with Neuropixels probes and software, J. Arnold, S. Sawtelle, P. Polidoro and M. Gugiu for help with setting up the maze, and Sandro Romani and Carsen Stringer for advice on analyses and comments on an earlier version of the manuscript.

## Author contributions

CB and AKL designed the study. CB performed the experiments and analyzed the data. CB and AKL wrote and edited the manuscript.

## Data and materials availability

All data required to reproduce the findings will be made available upon publication

## Supplementary Figures

**Figure S1.**
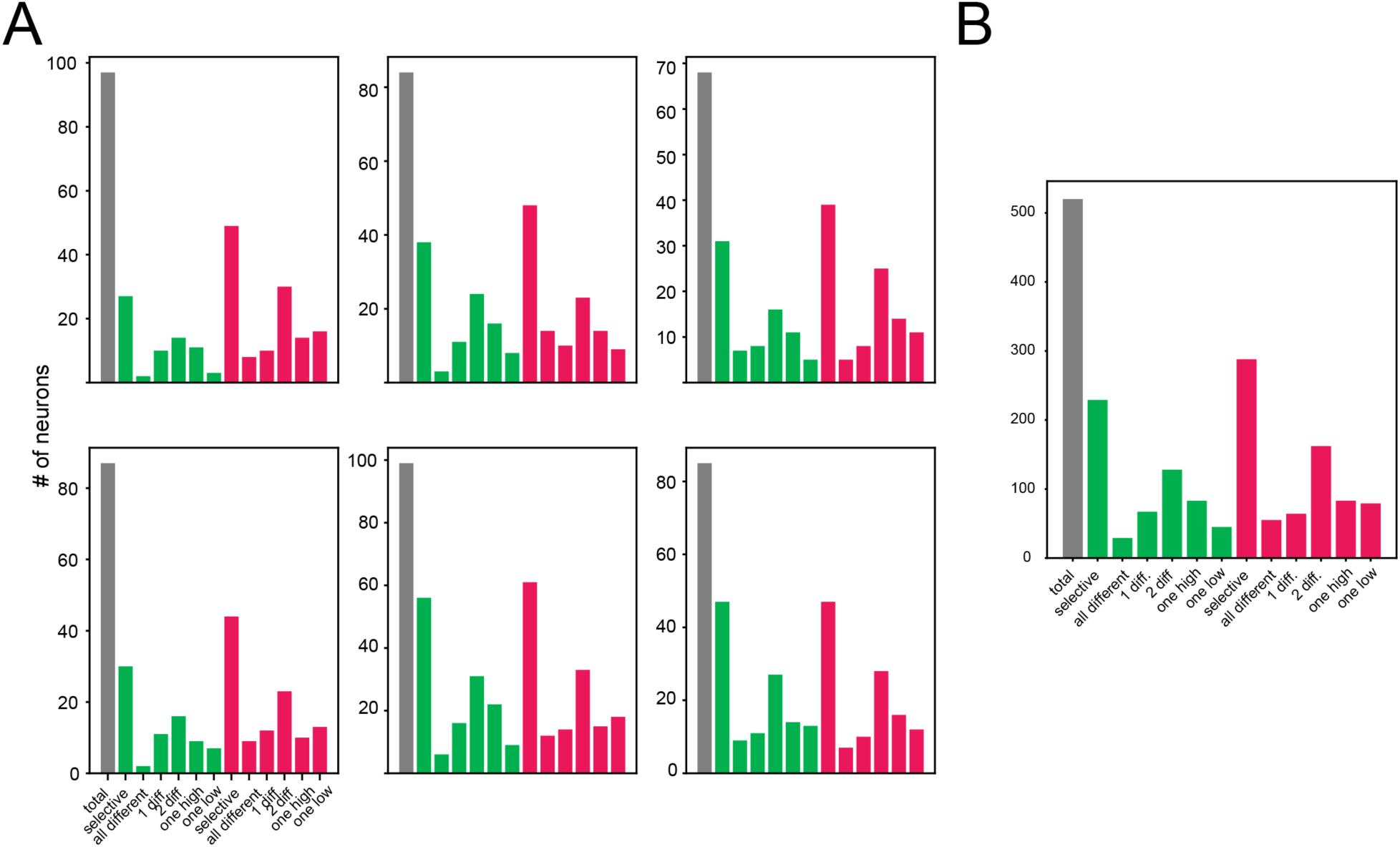
Types of spatial firing. Gray: total number of neurons per session. Green: at starts, red: at goals. Number of neurons with significantly different numbers of spikes at starts and goals (Kruskal-Wallis test). “All different”: number of neurons with differential firing at all three locations. “1 diff”: number of neurons where firing at one location is different from that at one other location. “2 diff”: number of neurons where firing at one location is different from that at the two other locations, and for these “2 diff” cells the firing rate at the location where it is different can be higher (“one high”) or lower (“one low”) than at the two other locations. **(A)** Single sessions. **(B)** All sessions pooled.

**Figure S2.**
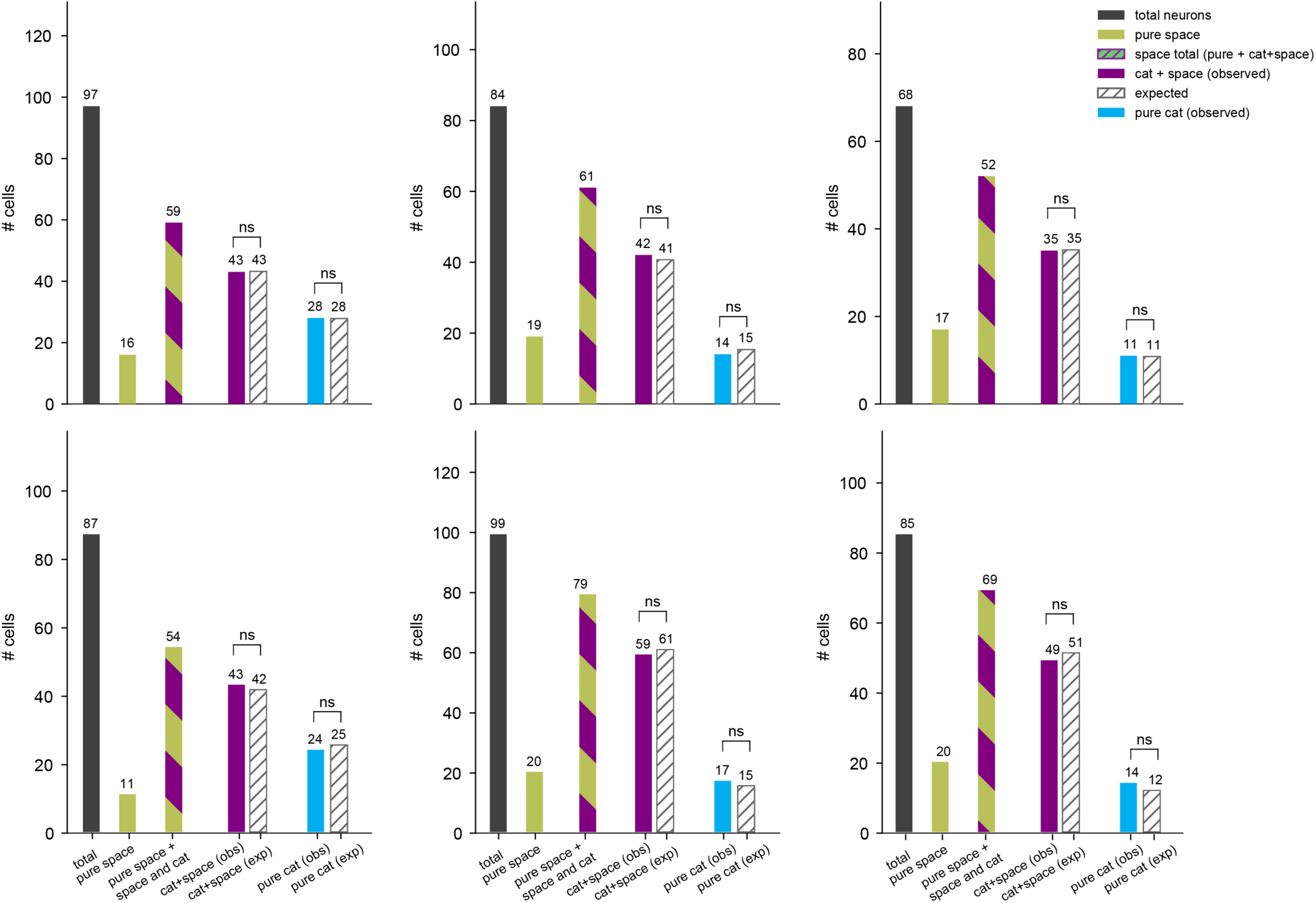
Distribution of category and spatial selectivity for single sessions. Compare to Fig. 2. Number of cells selective space only (‘pure space’), category only (‘pure cat’), and for various combinations. Number expected assuming selectivity for space and category are independent (hatched). Ns: p>0.05 (Fisher’s exact test).

**Figure S3.**
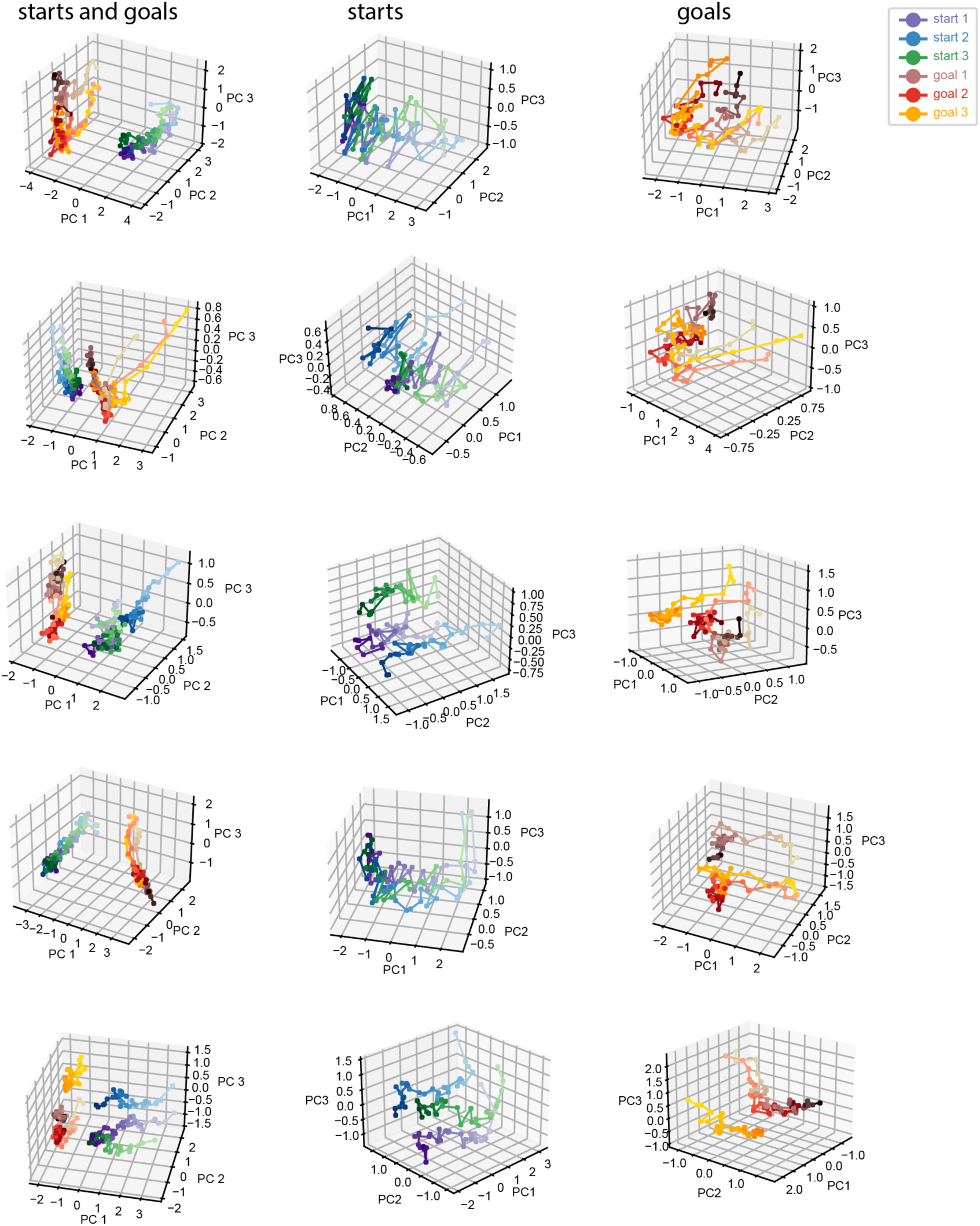
Principal component analysis for single sessions. Compare to Fig. 4 A, B. Left column: PCA of mPFC population activity (averaged over trials) at goals and starts, time bins 100 ms (and 106.7 ms for starts in 2 data sets). PCA for start (middle column) and at goal (right column) locations separately: note that time elapses in parallel for different locations and primarily along one principal component. Color shades from light to dark correspond to early to late time points at the starts/goals. Each row corresponds to one recording session.

**Figure S4.**
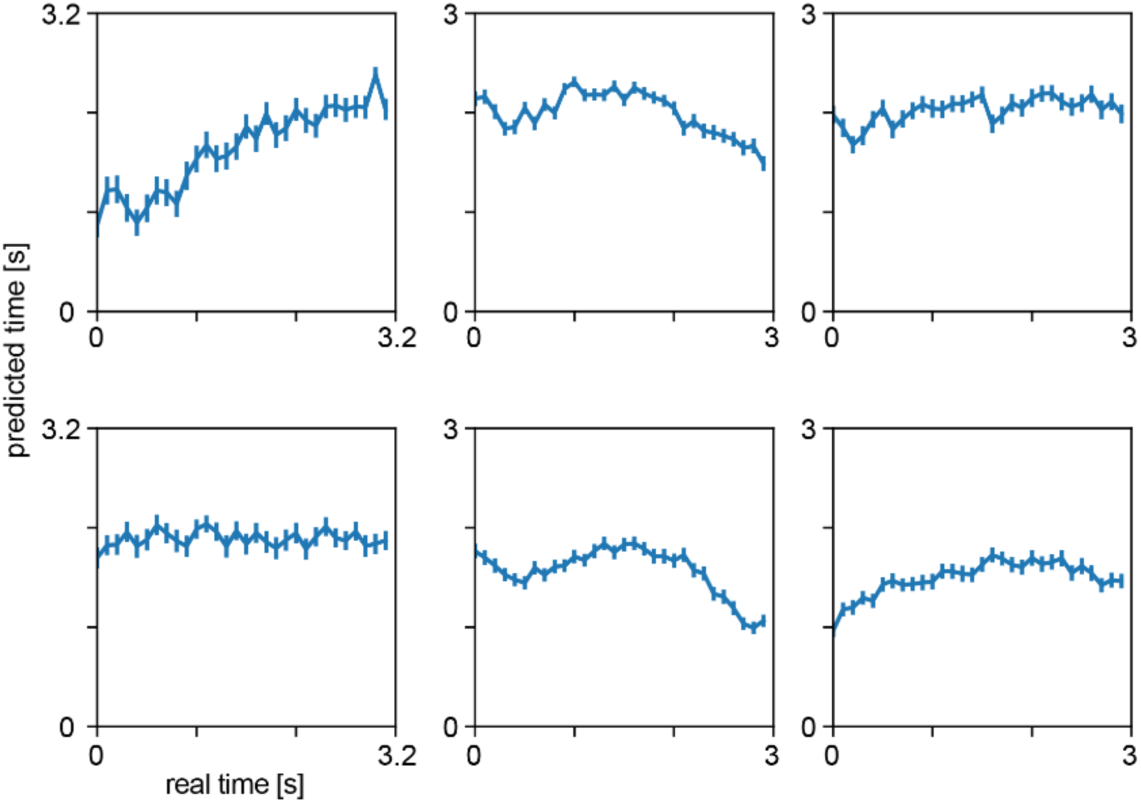
Time decoding across categories is inconsistent. For each session a logistic regression decoder was trained on start data and tested on goal data. Data at the start and at the goal were split into 30 time bins. Error bars denote standard error of the mean over trials. Each plot corresponds to one session. Different sessions show related, reversed, or no trend.

**Figure S5.**
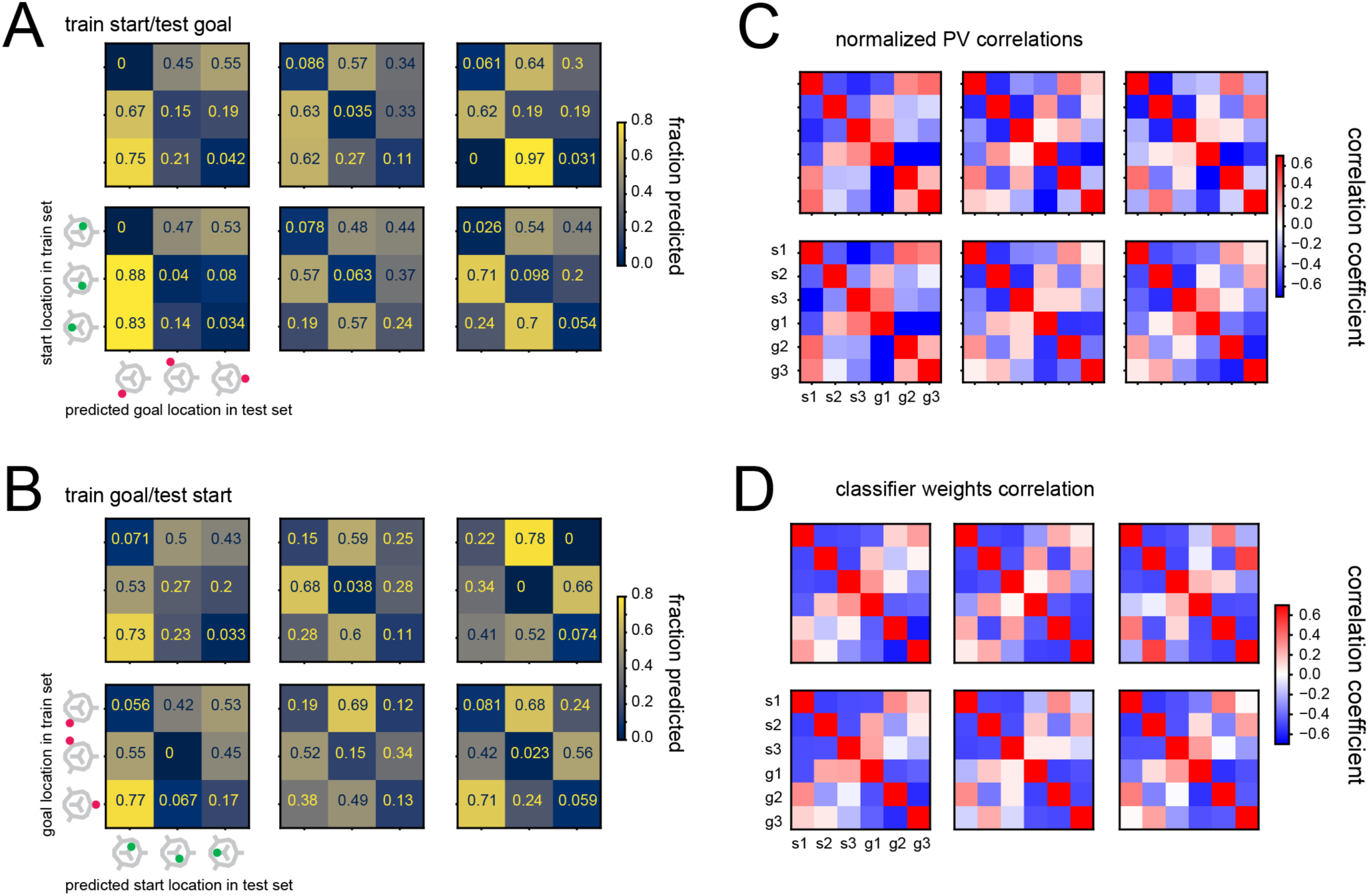
Start and goal locations have structured representations reflecting the geometry of the maze layout. **(A)** Predicting goal location with a classifier trained on start locations. Opposite locations (along the diagonal) were rarely predicted. **(B)** Predicting start location with a classifier trained on goal locations. Again, opposite locations were rarely predicted. **(C)** Correlation coefficients of population vectors of start and goal locations (the goal location and start location population vectors of all trials were standardized separately before correlation to reveal any structure beyond the high similarity among start and goal location population vectors). **(D)** Correlation of classifier weights obtained by separately training classifiers on start or goal locations. The six correlation matrices in each panel correspond to each of the six recording sessions. In C and D, the diagonals of the lower left and upper right 3×3 blocks in each matrix have low values, revealing that the activity at starts and goals opposite each other are least similar, as seen in A and B.

**Figure S6.**
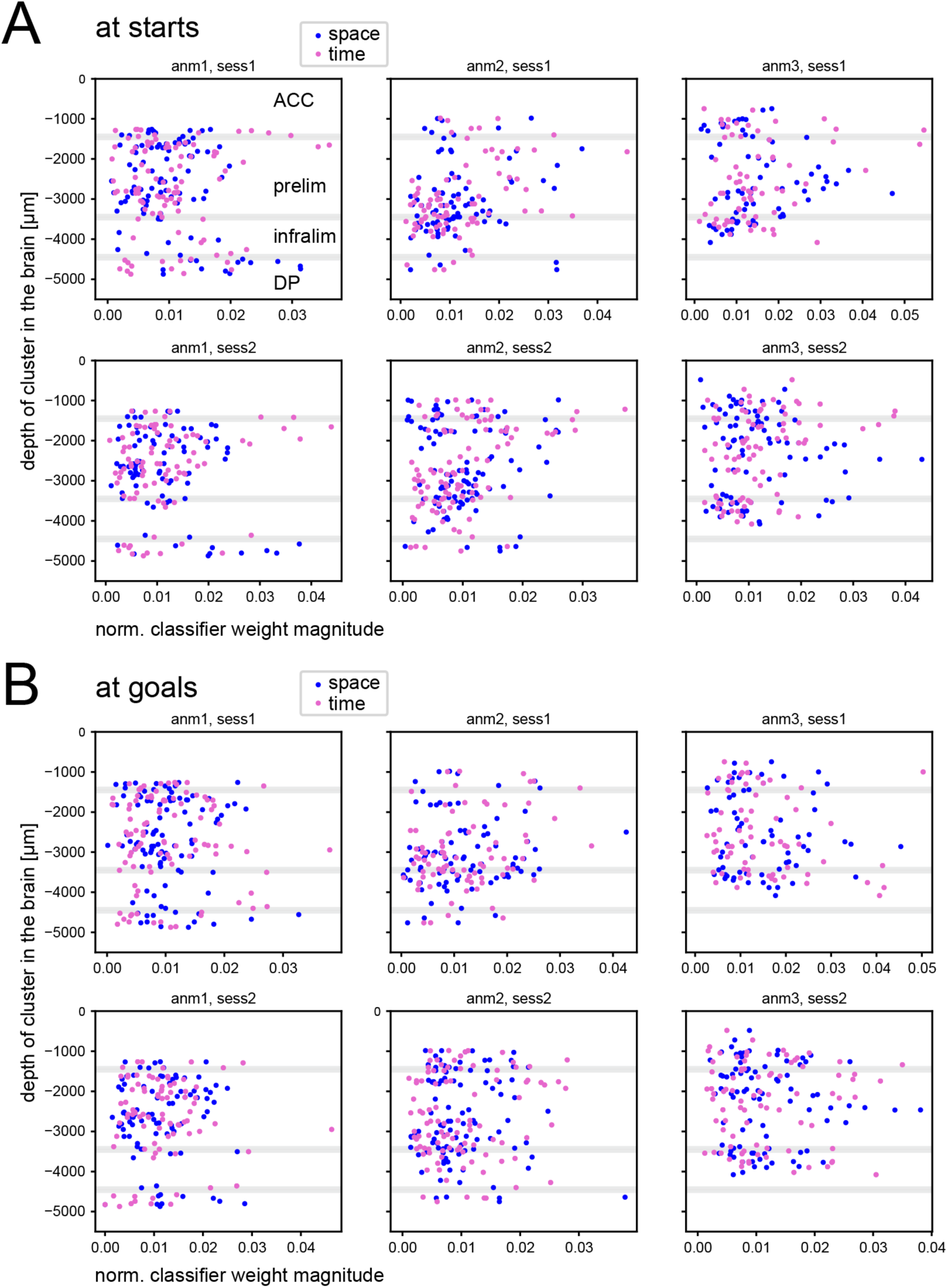
Anatomical distribution of space and time selectivity. Normalized classifier weights for all cells for space (blue) and time (pink) decoding versus depth of recording from the pial surface. Each scatterplot corresponds to one session. **(A)** At starts. **(B)** At goals. Divisions between subareas are estimates.

**Figure S7.**
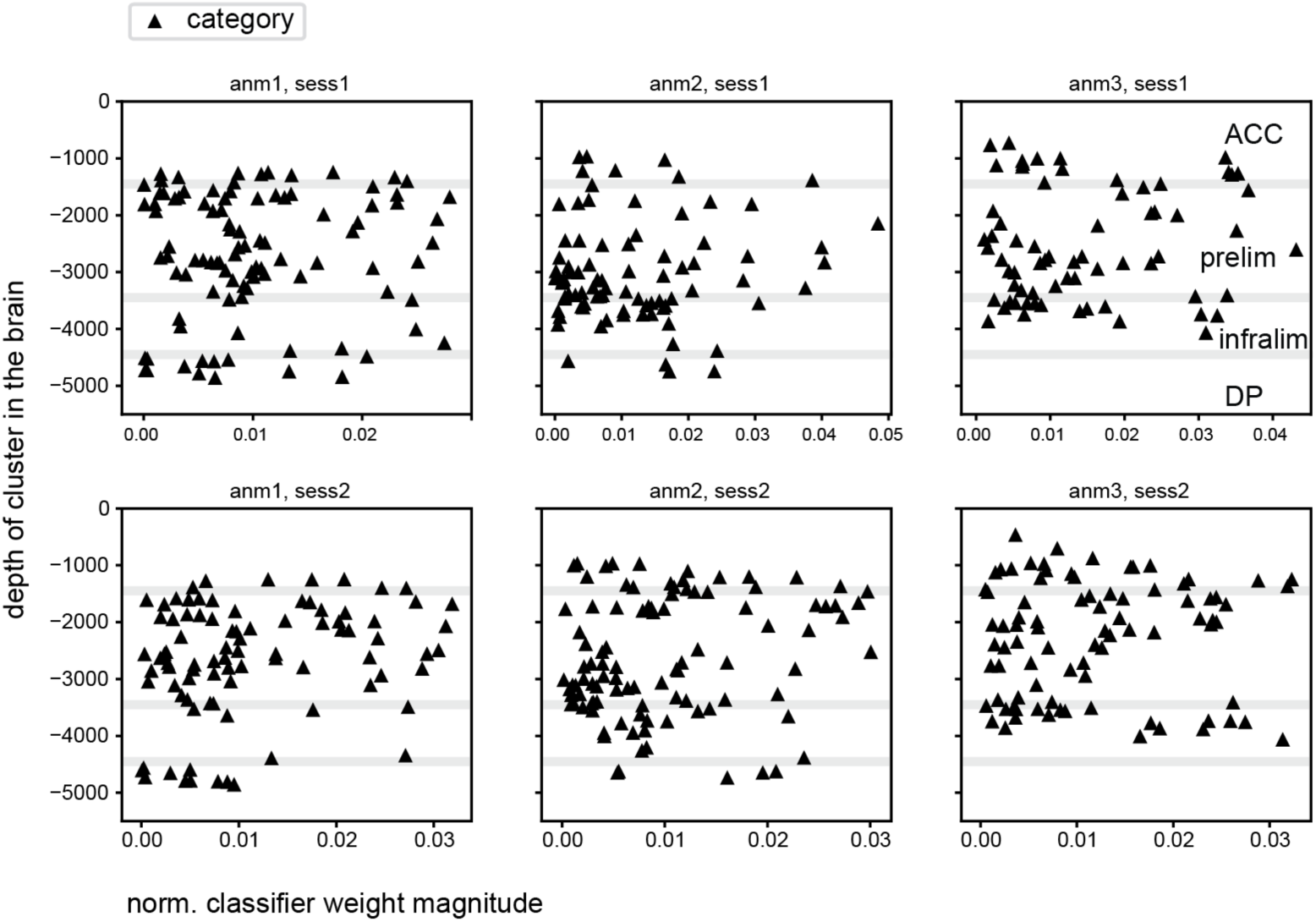
Anatomical distribution of category selectivity. Normalized classifier weights for all cells for category decoding versus depth of recording from the pial surface. Each scatterplot corresponds to one session. Divisions between subareas are estimates.

